# SwabExpress: An end-to-end protocol for extraction-free COVID-19 testing

**DOI:** 10.1101/2020.04.22.056283

**Authors:** Sanjay Srivatsan, Sarah Heidl, Brian Pfau, Beth K. Martin, Peter D. Han, Weizhi Zhong, Katrina van Raay, Evan McDermot, Jordan Opsahl, Luis Gamboa, Nahum Smith, Melissa Truong, Shari Cho, Kaitlyn A. Barrow, Lucille M. Rich, Jeremy Stone, Caitlin R. Wolf, Denise J. McCulloch, Ashley E. Kim, Elisabeth Brandstetter, Sarah L. Sohlberg, Misja Ilcisin, Rachel E. Geyer, Wei Chen, Jase Gehring, Seattle Flu Study Investigators, Sriram Kosuri, Trevor Bedford, Mark J. Rieder, Deborah A. Nickerson, Helen Y. Chu, Eric Q. Konnick, Jason S. Debley, Jay Shendure, Christina M. Lockwood, Lea M. Starita

**Author notes:** denotes equal contributors (Sanjay Srivatsan, Sarah Heidl, Brian Pfau).

## Abstract

**Background:** The urgent need for massively scaled clinical testing for SARS-CoV-2, along with global shortages of critical reagents and supplies, has necessitated development of streamlined laboratory testing protocols. Conventional nucleic acid testing for SARS-CoV-2 involves collection of a clinical specimen with a nasopharyngeal swab in transport medium, nucleic acid extraction, and quantitative reverse transcription PCR (RT-qPCR) (1). As testing has scaled across the world, the global supply chain has buckled, rendering testing reagents and materials scarce (2). To address shortages, we developed SwabExpress, an end-to-end protocol developed to employ mass produced anterior nares swabs and bypass the requirement for transport media and nucleic acid extraction.

**Methods:** We evaluated anterior nares swabs, transported dry and eluted in low-TE buffer as a direct-to-RT-qPCR alternative to extraction-dependent viral transport media. We validated our protocol of using heat treatment for viral activation and added a proteinase K digestion step to reduce amplification interference. We tested this protocol across archived and prospectively collected swab specimens to fine-tune test performance.

**Results:** After optimization, SwabExpress has a low limit of detection at 2-4 molecules/uL, 100% sensitivity, and 99.4% specificity when compared side-by-side with a traditional RT-qPCR protocol employing extraction. On real-world specimens, SwabExpress outperforms an automated extraction system while simultaneously reducing cost and hands-on time.

**Conclusion:** SwabExpress is a simplified workflow that facilitates scaled testing for COVID-19 without sacrificing test performance. It may serve as a template for the simplification of PCR-based clinical laboratory tests, particularly in times of critical shortages during pandemics.

## Introduction

Since the first reported cases in the winter of 2019, the spread of the novel beta-coronavirus SARS-CoV-2 has grown into a global pandemic. The virus spreads easily from person to person and is often carried by asymptomatic individuals (3). These viral properties, in conjunction with a lack of an effective centralized response or societal adherence to public health recommendations, has led to a continued persistence of the pandemic throughout the United States (4). It is widely recognized that increased testing capacity can ameliorate the outbreak (5,6), but the prohibitive cost of testing materials and reagents as well as global supply chain problems continue to thwart efforts to reach the required scale.

Since the beginning of the pandemic, the gold standard for SARS-CoV-2 detection has been extraction of purified RNA followed by reverse-transcription quantitative polymerase chain reaction (RT-qPCR). Specimens are traditionally collected as nasopharyngeal (NP) specimens (3) by healthcare professionals using a specific swab and transported in viral media (e.g. Universal Transport Media (UTM)). Worldwide reliance on this template protocol has led to global shortages in swabs, viral media, and laboratory reagents. These shortages continue to plague testing labs and impede efforts to scale. Prior literature (7,8) and the work of United Health/Quantigen (9), have established that swabs collected without transport media are acceptable for nucleic acid detection-based diagnostics, eliminating the reliance on UTM. Extraction-free protocols have also been developed to remove the need for RNA extraction reagents and streamline testing protocols. Saliva specimens have been shown to be particularly amenable to extraction-free testing protocols. For example, SalivaDirect – a protocol for performing SARS-CoV-2 RT-qPCR on saliva specimens without extraction (10) – had a sensitivity of 89% compared to traditionally processed anterior nares (AN) or oropharyngeal (OP) swabs, demonstrating the viability of extraction-free protocols. Unlike saliva, extraction-free methods for nasal swabs have been less sensitive than conventional protocols—likely due to PCR inhibition from transport media or saline (10–16).

Here we describe the development of an UTM and extraction-free protocol for anterior nasal dry-swabs that is compatible with RT-qPCR and does not sacrifice test performance. This protocol, which we have coined ‘SwabExpress’, has a low limit of detection, high sensitivity, high specificity, and superior test performance when compared to conventional extraction-based RT-qPCR protocols. We further identify and ameliorate two distinct failure modes for extraction free RT-qPCR-based testing. Widespread adoption of this approach and others like it could result in a dramatic increase in testing capacity, decrease consumables used during testing, and ultimately help curb the spread of SARS-CoV-2.

## Results

### Usability and Reliability of Anterior Nares (AN) Swabs for At-Home Specimen Collection

We first explored the use of anterior nares (AN) swabs for specimen collection. For mass testing purposes, a swab that is widely available, inexpensive, easy to manufacture, and simple for self-collection is critical. The US Cotton #3 swabs fit these specifications; a polyester AN swab that resembles consumer-brand Q-tips (17). For the purposes of scaled observed or at-home self specimen collection or specimen collection for a child, swabbing the anterior nares anatomical site would be more comfortable, accessible and easier to describe to test users leading to fewer mistakes and better specimen collection (18,19).

Therefore, we conducted a usability study to determine both the accuracy and ease of AN swabs in a Swab-and-Send program where at-home specimen collection kits were delivered to participant residences, the participants swabbed themselves or a child while being virtually monitored by clinical study coordinators and then packaged the specimen for return to the molecular testing lab (19,20). After using the specimen collection kit, study participants completed a survey reporting their level of confidence, the kit’s ease of use, and the level of discomfort experienced during swabbing. Participants were recruited from the greater Seattle area and spanned a range of ages, races, household income, and educational attainment (**fig. S1; Table S1A-D**).

The results of the usability study were very encouraging. The majority of participants reported only mild-discomfort during specimen collection with 40% of participants reporting no discomfort at all (**Fig. 1A**). A majority of study participants also found the instructions clear and felt confident that they had correctly collected their specimen (**Fig. 1B**). This was confirmed by low observed rates of error during specimen collection using the AN swabs and during packaging for return (**Table S2A-B**). Molecular testing performed on these self-collected specimens confirmed this; RT-qPCR detected the human marker RNase P gene in 100% of swabs with an average crossing threshold value of 23.5 (s.d. 1.7). The amount of RNase P recovered from the AN swabs was higher than for unsupervised collection of mid-turbinate swabs, which had an average crossing threshold (Ct) value of 26.9 (s.d. 2.5) (**Fig. 1C**). Together these data indicate that the use of widely available polyester swabs in the anterior nares is a viable and preferable alternative for at home specimen collection.

**Figure 1.**
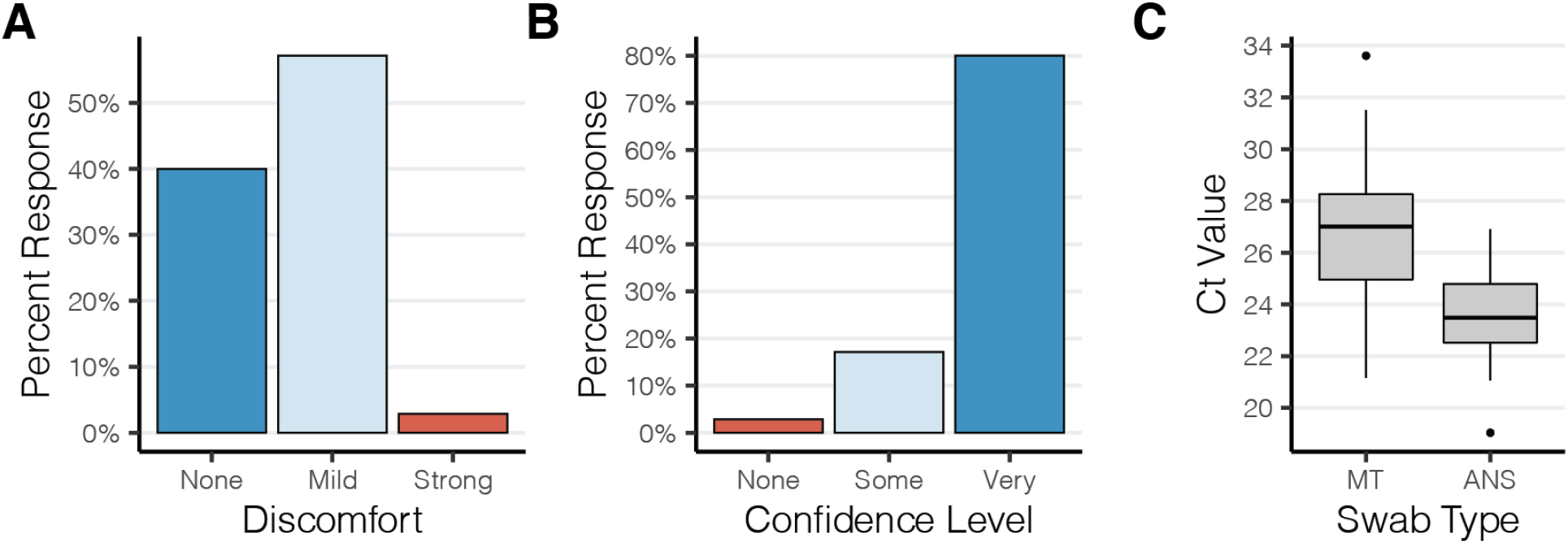
Polyester anterior nares swabs are both comfortable and easy to use. (A,B) Study participants’ (n=35) self reported **(A)** discomfort and **(B)** confidence during self administration of an anterior nares swab at home. **(C)** Boxplot depicting the RT-qPCR crossing threshold values for RNaseP from self-administered anterior nares swabs (ANS) and mid-turbinate (MT) swabs.

### Handling Dry Swabs in the Clinical Laboratory

Standard viral media such as Universal Transport Medium (*e.g.* COPAN Diagnostics) have been in short supply over the course of the pandemic. These salt-rich media inhibit direct RT-qPCR, making RNA extraction a necessity and thus create an additional bottleneck in the testing process. Furthermore, automated extraction systems are expensive and their reagents and consumables are also subject to global shortages. Therefore, we focused on eliminating UTM and extraction from our testing platform. To bypass UTM, we adopted a dry swab transport and rehydration method validated by Quantigen that has been explored by other clinical testing laboratories (15,21). Next, to eliminate RNA extraction and enable direct RT-qPCR, we tested rehydration solutions for their ability to elute contrived SARS-CoV-2 specimens, compatibility with direct RT-qPCR, and simplicity. We determined that elution in low-TE (10 mM Tris pH 7.5, 0.1 mM EDTA) without other detergents was best suited for direct RT-qPCR (**Fig. S2**). Unlike UTM and other saline solutions, the low ionic strength of low-TE does not inhibit PCR amplification. Moreover, low-TE can be quickly prepared using reagents commonly found in laboratories.

Bypassing nucleic acids extraction poses another problem; instead of the virus being inactivated by the denaturing agents during nucleic acid extraction, the specimen eluted from the swab remains potentially infectious for SARS-CoV-2 or other pathogens and poses a risk to laboratory staff. Accordingly, specimens from both conventional UTM and rehydrated dry swabs are processed inside a class II biosafety cabinet, in accordance with federal regulatory guidance. However, it is practical and beneficial for downstream steps (like preparing RT-qPCR reactions) to take place on a BSL-2 designated bench. Therefore, we compared several inactivation methods to determine which would be easiest without inhibiting PCR or causing a loss of sensitivity. Viral inactivation of coronaviruses can be achieved through the use of either detergent or heat (22). To test the viral inactivation of SARS-CoV-2 we compared both modalities to inactivate the virus. Heat inactivation at 65°C for 10 minutes or incubation in a buffer containing 0.25% Triton-X-100 at room temperature completely ablated the infectivity of high titre stocks of SARS-CoV-2. This was demonstrated using two complementary assays: 1. a lack of cytopathic effects was observed upon co-culture of inactivated virus with VeroE6 cells (a virus sensitive cell line) and 2. a lack of viral replication was observed by RT-qPCR when inactivated cultures were plated with Vero cells (a virus tolerant cell line) (**Fig. S3**). Taken with our prior results demonstrating the negative impact of detergents on RT-qPCR (**Fig. S2**), we opted to deploy heat inactivation. We used a protocol to heat inactivate at higher temperatures (95°C) for 30 minutes to increase the safety margins. We also determined that this high-heat protocol had the added benefit of stabilizing the sample over time, a result concordant with another SARS-CoV-2 testing protocol in saliva (23).

### Performance of Extraction-free RT-qPCR

Having developed an extraction-free RT-qPCR protocol (EF-RT-qPCR), we set out to determine its performance on both contrived and clinical specimens. To assess analytical sensitivity, we first determined this assay’s limit of detection (LoD); the minimum number of SARS-CoV-2 RNA molecules that could be detected in greater than 95% of RT-qPCR reactions. To generate these contrived specimens, we inoculated AN swabs with clinical matrix collected from a healthy volunteer with dilutions of heat inactivated SARS-CoV-2. These experiments determined the EF-RT-qPCR analytical sensitivity to be 2 molecules/μl of eluate for the Orf1b assay and 4 molecules/μl of eluate for the S-gene (Spike gene) assay (**Table S3**). This LoD is comparable to the LoD of many other RT-qPCR-based tests that have been issued Emergency Use Authorization from the FDA(24).

Next we tested the performance of EF-RT-qPCR compared to our clinically validated RT-qPCR laboratory-developed test on archived AN specimens. In this assay each sample is tested in 4 independent RT-qPCR reactions, comprising two SARS-CoV-2 assays (Orf1b and Spike) in duplicate, and is multiplexed with a RNase P assay in every well (**Fig. 2A** and **Fig S4**). Following RT-qPCR, a clinical result is determined by the number of replicates displaying SARS-CoV-2 amplification: positive (3 or 4 of 4 wells), low-positive/inconclusive (2 of 4 wells) and negative (0 or 1 of 4 wells) (**Fig. 2A,B**). Head-to-head comparison between EF-RT-qPCR and a reference standard extraction-based RT-qPCR assay on matched specimens established that EF-RT-qPCR was 100% specific (56/56 negative specimens) and 91.0% sensitive (61/67 - 56 positive and 5 low-positive) (**Fig. 2C; Table S4**). Comparison of the mean delta Ct (∆Ct) values between the two assays showed that eliminating extraction did decrease analytical sensitivity. We observed an average increase of 1.96, 2.45, and 4.00 cycles for Orf1b, Spike and RNase P assays, respectively. Indeed, the 6 specimens not detected by EF-RT-qPCR had an average Ct with the extraction-based RT-qPCR assay of 34.13 for Orf1b and 35.29 for Spike.

**Figure 2.**
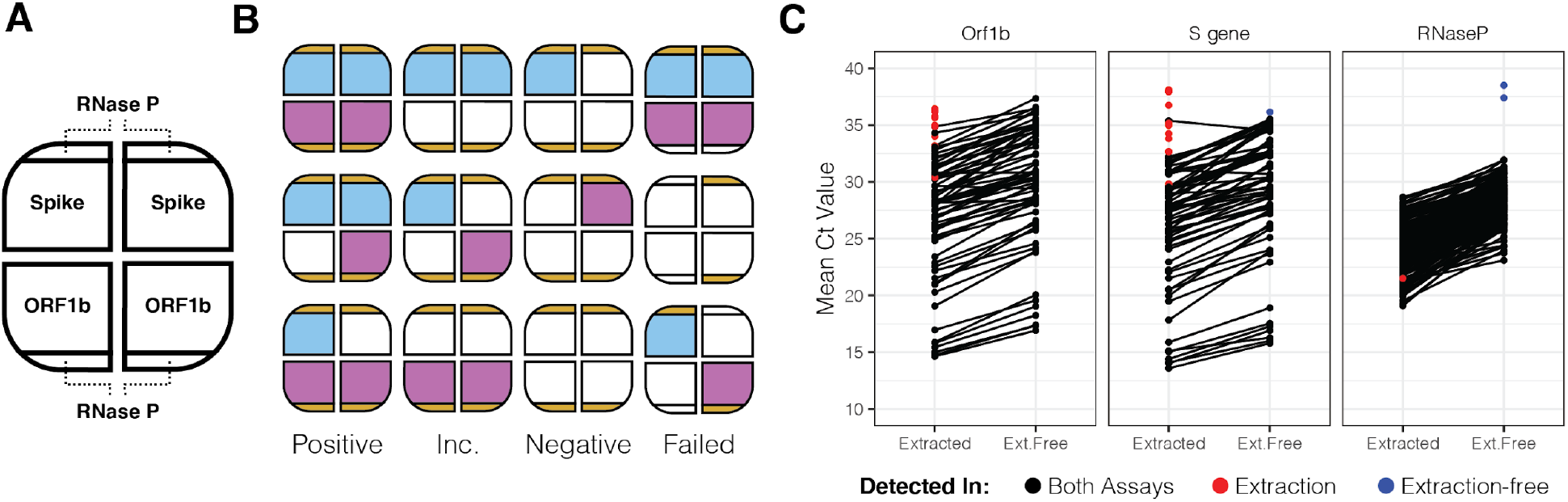
Extraction free RT-qPCR set-up and test performance. (**A**) Assay layout of the EF-RT-qPCR test. One sample is assayed in four wells on a 384 well-plate. Each sample is tested for two probes, in duplicate. RNase P is assayed in each well. (**B**) Combinations of how positive, inconclusive (abbreviated Inc.), negative, and failed samples are determined. (**C**) Mean Ct values for 67 specimens processed by EF-RT-qPCR and extraction-based RT-qPCR.

Owing to an unstable supply chain, while validating the EF-RT-qPCR protocol, our clinical laboratory was forced to switch from the Roche Magna Pure 96 to the Thermo Fisher KingFisher Flex automated nucleic acids extraction platform. The relative sensitivity, specificity and ∆Ct values between 619 prospective specimens run in parallel on both the KingFisher Flex (extraction) and EF-RT-qPCR were comparable to results of the retrospective study on stored specimens. EF-RT-qPCR detected SARS-CoV-2 in 100% of specimens that were positive by the extraction method with a 99.4% specificity (**Tables S5–S7**).

### Addition of Proteinase K reduces amplification interference

After deploying EF-RT-qPCR as our clinical testing platform, we repeatedly observed two undesirable outcomes that were not observed in our validation studies. First, for 0.9% of specimens (n = 383/43,539), amplification of the human RNase P internal control was undetected in 2 or more of the 4 reactions (**Fig. 2B**; **Table S8**). These specimens were classified as “failures” and each sample was repeated before releasing the result. Second, for 0.5% of specimens (229/43,539), we sporadically observed the presence of strong amplification (Ct < 30) in a single well for one of the SARS-CoV-2 targets in specimens where the three other wells were undetected (**Table S9)**. However, upon repeat RT-qPCR, both with and without extraction, all 4 wells of the SARS-CoV-2 reactions for these specimens were undetected.

We noted that some of the specimens that produced these problematic outcomes had excess mucous or other nasal secretions. Therefore, we hypothesized that the addition of proteinase K (ProK) digestion could ameliorate both RNase P failures and the spurious SARS-CoV-2 amplification by digesting mucins and other potentially interfering proteins in the nasal specimens (25). We compared RT-qPCR results for 1,222 clinical specimens prepared by the 30 minute 95°C heat treatment with those digested with ProK for 15 minutes prior to heat treatment at 95°C for 15 minutes. We observed approximately 10-fold fewer RT-qPCR reactions with failed RNase P amplification -- 27 of 4,888 without ProK vs 2 of 4,888 with ProK -- reducing the repeat rate to 0.04% (**Fig. 3A, Table S10**), and improved RNase P detection (∆Ct −0.88) (**Fig. S5**). Furthermore, the addition of a ProK digestion step eliminated spurious amplification of SARS-CoV-2 targets.

**Figure 3.**
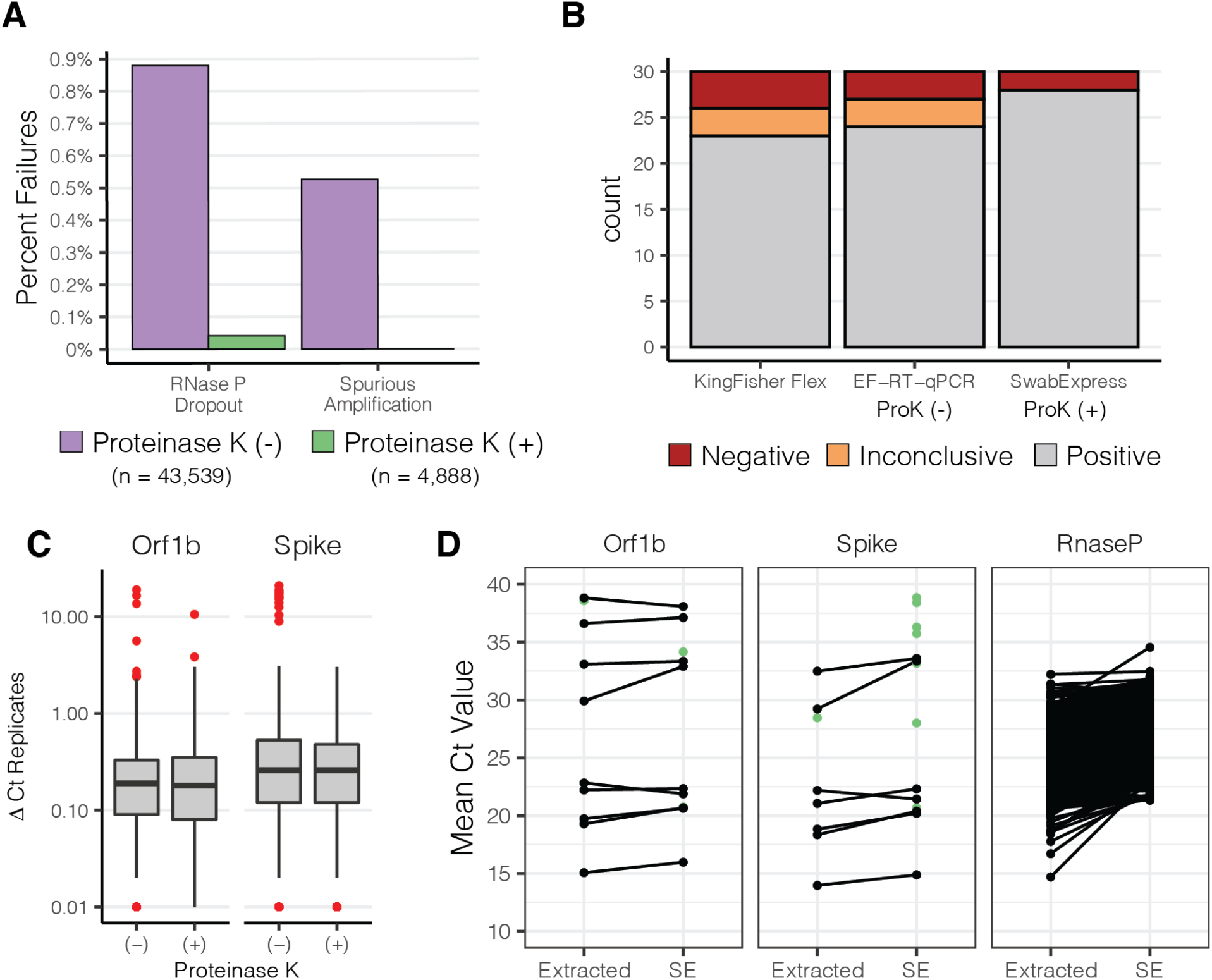
Addition of Proteinase K improves test performance. **(A)** Observed percentage of test failures with and without the addition of Proteinase K. **(B)** Archived samples with Ct > 28 reprocessed with either KingFisher Flex Extraction (left), Extraction-Free RT-PCR (middle), or SwabExpress (Extraction-Free RT-PCR + ProK) (right). Colors signify the number of samples and their classifications. **(C)** Box and whisker plots depicting the average delta Ct between replicate wells for SARS-Cov2 positive specimens. Red points indicate outliers. ∆Ct values were more consistent upon addition of Proteinase K. **(D)** Mean Ct values of matched specimens run through the automated KingFisher extraction system (left) or using SwabExpress (SE). Specimens which were detected in only one of the two protocols are displayed as green points.

In the 4,888 specimens processed both with and without ProK, ProK-treated specimens had decreased Ct values (mean decrease of 1.22 for Orf1B, and 0.97 for Spike). This increased sensitivity was also reflected in the ability to accurately classify archived SARS-CoV-2 positive specimens with Ct values > 28 (**Fig. 3B**). Repeatability and reproducibility were also improved with the addition of ProK (**Fig. 3C**). Upon addition of ProK, on SARS-CoV-2 positive samples , our protocol had a higher concordance (93.3%) versus without ProK (90%) or specimens extracted on the KingFisher Flex (86.6%) (**Table S11**). After this optimization we named our final protocol “SwabExpress” -- consisting of a dry AN swab, followed by ProK digestion and direct RT-PCR. Finally, we prospectively compared performance on 1169 specimens run in parallel on the SwabExpress and KingFisher Flex (extraction) platforms. Positive and negative clinical concordance was excellent; there was 100% concordance on positives calls, 99.91% concordance across negatives with a small ∆Ct value of 0.37 for the Orf1b target and 1.46 for the S target between the two assays (**Fig. 3D**, **Table S12**).

### SwabExpress is compatible with other SARS-CoV-2 RT-qPCR assays

Our laboratory-developed test uses custom Orf1b and Spike-gene assays for detecting SARS-CoV-2. To establish that the SwabExpress protocol was compatible with the widely used CDC N1 and N2 assays, we performed RT-qPCR on 75 positive specimens and 92 negative specimens with the N1 and N2 assays run in parallel on the SwabExpress platform and extraction-based RT-qPCR platform. The results were 100% concordant between our custom assays and the CDC assays. Ct values for positive samples were delayed when prepared by SwabExpress protocol compared to the Roche Magna Pure 96. However, this difference did not change the clinical interpretation of these samples (**fig. S6**). For the N1 assay, extracted specimens had an average Ct of 19.22 ± 3.67 versus 21.79 ± 4.33 with SwabExpress (∆Ct of 2.57). For the N2 assay, extracted specimens had an average Ct of 18.31 ± 3.73 versus Cts of 19.80 ± 3.72 for SwabExpress + proteinase K protocol (∆Ct of 1.49) (**Table S13).**

### SwabExpress is time and cost effective

A dry-swab, extraction-free RT-qPCR protocol comprises the minimal components of a diagnostic test. Although the addition of a proteinase K digestion adds $0.14 USD to the reagent cost for each sample, this cost is warranted. The addition of proteinase K reduces the repeat rate, reduces the chances of a false positive result from interfering substances during PCR amplification, improves the performance of the test, and in our hands, outperformed a suboptimal yet widely used automated extraction system (Thermo KingFisher Flex™).

Upon adoption, SwabExpress approximately doubled laboratory capacity. First, hands on technician time previously spent preparing and running extraction systems, went towards accessioning and processing additional samples. Second, the SwabExpress protocol increases scale by using a convection oven that can process upto 6 96-well plates simultaneously. This throughput greatly exceeds the single 96-well plate processed by commercial automated extraction systems. Further scaling of the SwabExpress protocol can be accomplished through the purchase of additional or larger ovens, although RT-qPCR instruments used during amplification and readout, still pose a substantial bottleneck in the testing protocol.

Along with the substantial cost of purchasing automated extractors, the consumables required for their operation cost between $4 and $5 per sample. By eliminating extraction, and transport medium, SwabExpress reduces the associated costs by more than 90% (~$0.20 per sample). In all, SwabExpress offers a time and cost-saving alternative to nucleic acids extraction using readily available reagents, which reduces dependence on a heavily burdened supply chain (**Fig. 4, Table S14**).

**Figure 4.**
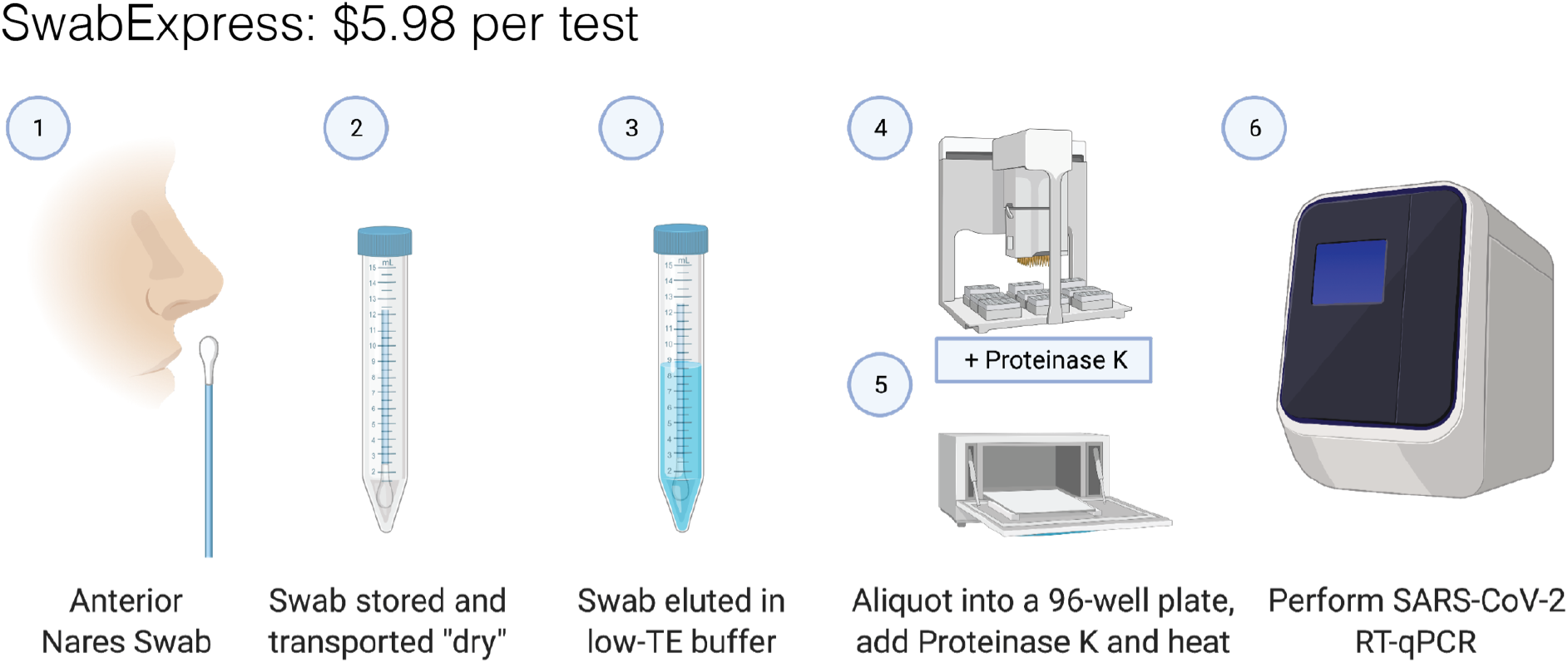
SwabExpress workflow. **(1)** Anterior nares swabs are collected and (2) transported dry to the lab. Upon receipt, (3) each swab is then hydrated with low-TE buffer, aliquoted into a 96 well plate, and (4) proteinase K is added to every well. (5) The eluted specimens are digested and heat-inactivated in a laboratory oven before (6) they are loaded as the template in a RT-qPCR reaction. The cost listed includes reagents and consumables.

## Discussion

Here we present SwabExpress, an end-to-end diagnostic platform optimized for faster and simpler low-cost detection of SARS-CoV-2 from nasal swabs without the use of nucleic acid extraction (**Figure 4)**. This protocol was so named for its ease, rapid turn around and simplicity -- dry swabs, without extraction, enhanced with proK digestion. By eliminating transport media and extraction from the workflow, we have decreased cost per sample and reduced supply chain pressure for the lab. Because of the reduced cost and the ability to process many more specimens in parallel, our lab’s capacity markedly increased with its adoption. Importantly, we gained efficiency without sacrificing accuracy; our results suggest that the simplified SwabExpress protocol (direct elution from dry swab into low-TE + proteinase K → RT-qPCR) is as sensitive as the conventional PCR protocol (swab → UTM → RNA extraction → RT-qPCR). SwabExpress has supported scaled testing in our lab with over 72,000 tests performed to date and allowed us to support large testing endeavors such as the Husky Coronavirus Testing Program for the University of Washington (26).

There are some caveats to consider. Even with the addition of proteinase K, specimens with excess mucous fail to amplify RNase P. Since adding this proteinase digestion step, 18/12,991 specimens have had two or more RNase P reactions fail (0.1%) and our laboratory reflexes these few specimens to an extraction protocol. However, 0.1% compares favorably when compared to a protocol with extraction where the failure rate due to failed RNase P is 1% (215/22,546). In addition, the unknown presence of inhibitors precludes comparison of Ct values between specimens; therefore, studies directly comparing Ct values from different specimens may not yield accurate results.

We have observed a marked loss of viral RNA after freeze-thaw cycles for specimens stored in low-TE compared to specimens stored in commercial UTM. We detect a ∆Ct of ~2.5 for specimens after −80℃ storage in low-TE, whereas the ∆Ct for specimens retested after storage at −80℃ in UTM has historically been negligible. This impacts the ability to use these specimens for downstream applications such as genomic sequencing.

Several improvements can be incorporated into the SwabExpress platform. First, the ability to detect multiple pathogens from one assay can be explored. It is likely that SwabExpress will be compatible with other enveloped viruses such as influenza and respiratory syncytial virus. Multiple targets can be detected in many qPCR systems and the reemergence of these viruses as COVID-19 prevention control measures begin to relax will be of interest to monitor. Second, the most labor intensive part of the SwabExpress protocol is sample accessioning. Receiving individual 10 ml tubes and transferring the eluate to 96-well format takes ~2.5 minutes per sample. Receiving nasal swabs in 96-well compatible, lab-ready transport tubes would streamline the process considerably.

Massive scaling and deployment of SARS-CoV-2 testing is essential to curtailing the COVID-19 pandemic, and will likely be necessary well into the future. The protocol evaluated here, including thousands of real-world, self-collected nasal swabs, would markedly simplify the workflow for RT-qPCR, the most widely deployed testing paradigm, by eliminating the need for viral transport media and RNA extraction, both of which are currently experiencing significant supply chain challenges. Looking forward, we envision that nasal swabs -- self-collected into laboratory ready, barcoded tubes and transported dry -- could potentially serve as a common input to a range of SARS-CoV-2 nucleic acid tests for public health surveillance applications. This includes gold-standard tests like RT-qPCR, but also potentially new modalities like Swab-Seq (*11*). The operationalization of the mass distribution and return of such lab-ready collection devices is a significant effort that should begin now.

## Methods

### Collection of Nasal Swabs

For preliminary studies (summarized in **Fig. S2**), individuals who tested positive for SARS-CoV-2 through routine clinical testing were identified and recruited into a study of home-based, self-collected home swabs (27). After providing consent, enrolled participants were supplied a swab and send kit (19) containing two swabs (Copan FloqSwab 56380CS01) delivered to their home via 2-hour delivery and were provided instructions to self collect two mid-turbinate swabs. Participants placed one swab in a tube with UTM (Becton Dickinson PN 220220) and the other in an empty, dry 15 mL conical tube for transport. For all other studies, anterior nares (US Cotton #3, distributed by Steripack) swabs were collected by the Seattle Flu Study, Husky Coronavirus Testing Program (HCT) (26) or the Seattle Coronavirus Assessment Network (SCAN) (28). Anterior Nares swabs were transported in a sterile, empty conical tube directly to the lab by HCT technicians. SCAN swabs were packaged by the participant according to kit instructions and sent to the Brotman Baty Institute / Northwest Genomics Center, utilizing standard IATA shipping procedures by courier at ambient temperature. These IRB-supervised studies were public health surveillance programs and enrolled both symptomatic and asymptomatic participants. Informed consent was obtained from adult participants and parents/permanent legal guardians of participant children. Archived and fresh convenience specimens from these studies were chosen at random for these studies.

### Usability Study

To recruit a sufficient number of children for the prospective usability study, participants were recruited that met broad eligibility criteria: 1. No COVID-19 symptoms, 2. no prior self-swab experience and 3. no prior medical or laboratory training. We obtained informed consent from adult participants and parents/permanent legal guardians of participant children.

### Swab rehydration and elution

All work was performed within a class II biosafety cabinet with appropriate precautions. For preliminary studies (**Fig.S2**) each collected mid turbinate dry swab was first removed from its 15 mL conical tube and placed into a 1.5 mL microfuge tube. Swabs were then cut using a sterile razor blade. Next, 200 μL of Tris-EDTA [10mM Tris-HCl pH 7.5 (T2319-1L, Sigma), 0.1mM EDTA (15575020, Invitrogen)] was added to each tube and vortexed for 30 seconds. Microfuge tubes containing swabs were then placed in a microcentrifuge and spun for 5-10 seconds to collect eluate. To test various buffers, 45 μL of this solution was removed and added to either 5 μL of low TE or 5 μL of 10% Triton-X (X100-500ML, Sigma Aldrich). These two specimens constitute the undiluted eluate from the dry swabs.

For all other studies anterior nares swabs were rehydrated in 1 mL low-TE (10 mM Tris pH 7.5, 0.1 mM EDTA) prepared in sterile, UltraPure DNase/RNase-Free Distilled Water (Life Technologies PN 10977023). Specimens were vortexed for 30 seconds or shaken for 1 minute and allowed to incubate at room temperature for at least 10 minutes before transfer to Matrix tubes (Thermo Fisher).

### RNA extraction of specimens

200 μL of eluate was extracted on the Magna Pure 96 using a DNA and Viral NA Small Volume Kit (Roche, 06543588001) with the universal small volume protocol and eluted into 50 μL proprietary elution buffer. After October 18, 2020, 200 μL of eluted anterior nares specimens were extracted on the KingFisher Flex using the MagMAX Viral Pathogen II Nucleic Acid Isolation Kit with MagMAX™ Viral/Pathogen Ultra Enzyme Mix (Thermo Fisher A48383 and A42366) and eluted in 50 μL (although ~35 μL is eluted).

### SwabExpress Specimen Preparation

50 μL of 94 specimens in Matrix tubes were transferred to a LoBind 96 well plate (Eppendorf 30129512) using a manual 96-well pipetting system (Rainin Liquidator) with low retention tips (Rainin 17014402) with or without 5 μL of Proteinase K (Thermo Fisher A42363) preloaded in the plate. The plate was sealed using an Eppendorf heat sealing foil (Eppendorf 0030127854) and an Eppendorf HeatSealer S200 (Eppendorf 5392000013) at 160°C settings for 4.5 seconds. Specimens with Proteinase K were incubated at 37°C for 15 minutes in a forced air convection oven (Across International 0853924003042) and then transferred to a second oven for heat inactivation at 95°C for 15 minutes. Specimens without Proteinase K were heat inactivated at 95°C for 30 minutes. The full standard operating procedure for SwabExpress is included as a supplemental file.

### RT-qPCR

Each RT-qPCR reaction was performed at a final volume of 10 μL and containing 1X TaqPath RT-qPCR MasterMix (PN A15300, Life Technologies), RNAse P TaqMan VIC assay (A30064, Life Technologies) or RNAse P HEX assay (IDT), SARS-Cov-2 ORF1b FAM assay (PN 4332079, Life Technologies assay# APGZJKF) or spike (S) gene (PN 4332079, Life Technologies assay# APXGVC4) and nuclease-free water (1907076, Thermo Fisher). Primer sequences were designed against Wuhan-Hu-1 sequence #MN908947.3 and are proprietary to Thermo Fisher. After dispensing 5 μL of these master mixes to each well of a 384 well plate (Applied Biosystems PN 4309849) using a Mantis Liquid Handler (Formulatrix), 5 μL of extracted or heat treated specimen was added to each well. Plates are sealed using optically clear microseal B (Biorad). Each assay is performed in technical duplicate for a total of four RT-qPCR wells per sample (**fig. S3**). RT-qPCR was then performed on the Applied Biosystems QuantStudio 6 Pro (25°C for 2 minutes, 50°C for 15 minutes, 98°C for 3 minutes, followed by 40 cycles of 98°C for 3 seconds and 60°C for 30 seconds). Reported Ct values were obtained from the onboard analysis using predetermined cycle thresholds. Positive controls contained purified nucleic acid with sequence that was amplified by the ORF1b and Spike gene assays.

The RT-qPCR reaction for the CDC COVID-19 diagnostic test was performed at a final volume of 20 μL. Reactions contained 1X TaqPath RT-qPCR MasterMix (PN A15300, Life Technologies), nCOV-N1 FAM or nCOV-N2 FAM primer and probe mix (10006713, IDT) and nuclease-free water (1907076, Thermo Fisher) and 5 μL of sample prepared by extraction or SwabExpress was added to each well. RT-qPCR was then performed on the QuantStudio 6 Pro (25°C for 2 minutes, 50°C for 15 minutes, 98°C for 3 minutes, followed by 40 cycles of 98°C for 3 seconds and 60°C for 30 seconds). Reported Ct values were obtained from the onboard analysis using the auto-determined cycle thresholds. Data was analyzed using Excel and R 3.5.

### Preparation of inactivated viral controls

For contrived SARS-CoV-2 positive mid turbinate swab specimens, two healthy volunteer self-swabs were administered and collected for preliminary studies (**Fig. S2**). Each swab was then loaded with 2 μL of heat-inactivated virion (VR-1986HK [1.6e6 virion/μL], ATCC) each dry swab was allowed to dry to completion for 6 hours at room temperature in an uncapped 15 mL conical tube. For the anterior nares swabs, contrived SARS-CoV-2 positive swabs were generated by collecting clinical matrix from a confirmed healthy volunteer and loaded with 2 μL of diluted heat-inactivated virion (VR-1986HK [1.6e6 virion/μL], ATCC).

### Viral Inactivation Studies

Viral inactivation studies were performed at Seattle Children’s Research Institute’s biosafety level 3 facility. Twenty-five μL of SARS-CoV-2 (isolate USA-WA1/2020 obtained from ATCC BEI Resources) viral stock with a titer of 5.8×10^6^ pfu/mL was incubated in 200 μL of TE or TE + 0.25% Triton for 10 minutes at room temperature, or in TE at 65°C for 10 minutes. Untreated and treated SARS-CoV-2 was then added neat and at 10-fold dilutions through 10^−7^ to confluent cultures of Vero E6 cells (CRL-1586, ATCC) and 48 hours later cytopathic effects were scored after staining with crystal violet. RNA was isolated from Vero cells using a TRIzol Plus RNA Purification Kit (ThermoFisher) and the amount of SARS-CoV-2 was quantified by RT-qPCR.

### Retrospective Comparison Studies Between Testing Methods

Remnant participant specimens were stored either at 4°C or −80°C and prepared by extraction or heat treatment or heat treatment + proteinase K digestion as described above. Technicians performing testing and clinical directors interpreting results were both blinded to previous test results.

### Prospective Comparison Studies Between Testing Methods

freshly acquired specimens from the SCAN and HCT studies were prepared by extraction or heat treatment or heat treatment + proteinase K digestion and tested by RT-qPCR in parallel. For prospective analyses, both technicians and clinical directors performed testing and interpretation blinded to results from the comparator method.

### Ethics Approval

Sequencing and analysis of specimens from the Seattle Flu Study, the Hospitalized and Ambulatory Adults with Respiratory Viral Infections (HAARVI) study and the SCAN study were approved by the Institutional Review Board at the University of Washington (protocols STUDY00006181, STUDY00000959, STUDY00007628, STUDY00010432, STUDY00011148). Informed consent was obtained for all participant specimens.

## Acknowledgements

We would like to thank the Seattle Flu Study, HAARVI, SCAN and Husky Coronavirus Testing program study participants for their invaluable contributions to this research. We thank Sydney Floth, Jefferson Nguyen, Ashley Gate, and Gift Nwame for help with the usability study, Catherine Moore of Public Health Wales and Dan Wattendorf, Emily Turner and Karen Heichman of the Bill and Melinda Gates Foundation for helpful advice.

The Seattle Flu Study and SCAN are administered by the Brotman Baty Institute for Precision Medicine and funded by Gates Ventures, the private office of Bill Gates. The funder was not involved in the design of the study and does not have any ownership over the management and conduct of the study, the data, or the rights to publish. Funding for the HAARVI study comes from DARPA HR001117S0019, the Bill and Melinda Gates Foundation, Emergent Ventures grant to HYC. LMS and JS are funded by 1RM1HG010461-01 from the NHGRI and JS is an Investigator of the Howard Hughes Medical Institute. JSD is funded by NIH/NIAID K24AI150991-01S1, which supported the viral inactivation studies. Additional funding for this project came from the University of Washington in support of the Husky Coronavirus Testing Program with funds from the United States Senate and House of Representative Bill 748, Coronavirus Aid, Relief, and Economic Security Act (CARES Act).

---

**Supplementary Figure 1.**
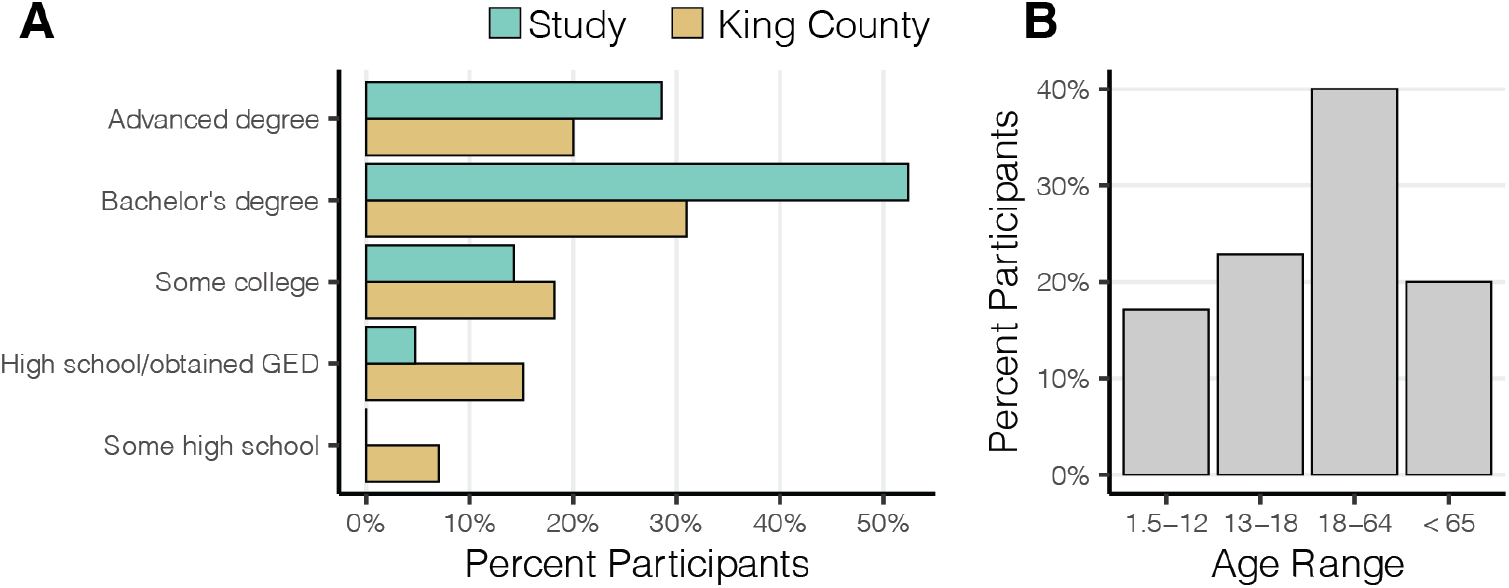
Educational attainment and age range of 35 swab usability study participants. (**A**) The highest level of education attained by recruited participants (teal) versus King County, Washington (golden). (**B**) Percentage of recruited study participants in the displayed age ranges.

**Supplementary Figure 2.**
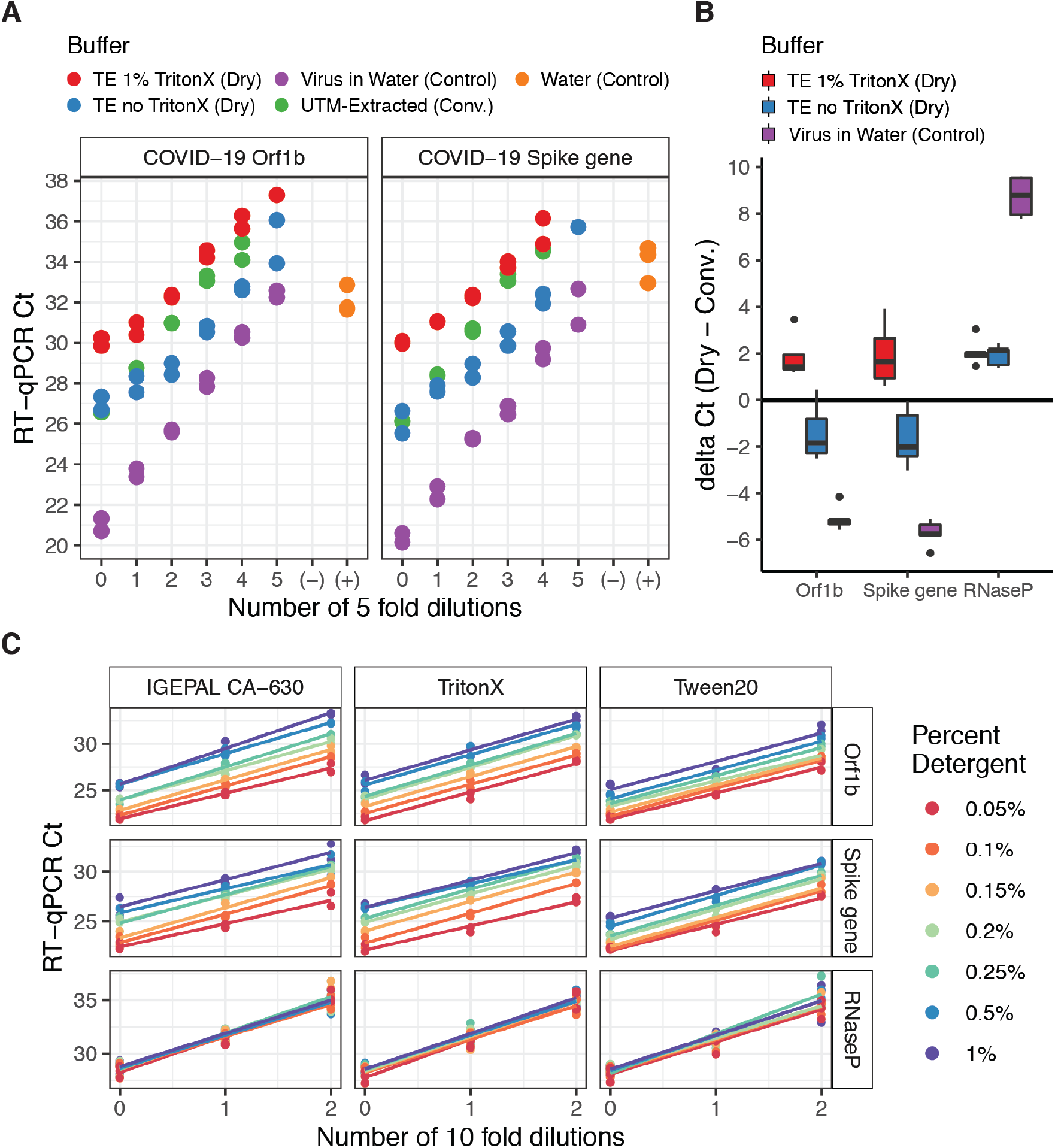
Comparison of RT-qPCR detection of inactivated virus from conventional and dry swabs. (**A**) Crossing threshold (Ct) values shown for specimens comprising a self swab and inactivated SARS-CoV-2 virus for both the ORF1b primer-probe set (left) and the Spike gene primer-probe set (right). Colors correspond to unique combinations of extraction protocol or controls. All specimens were measured twice in independent RT-qPCR reactions. No template control (−) wells contained either buffer or water and positive control wells (+) contained synthetic template. (**B**) Delta Ct values between conventionally processed swabs and dry processed swabs at matched dilutions for this contrived experiment. (**C**) Ct values for three probes (rows: Orf1b, Spike, Rnase P) assayed in buffers containing one of three detergents (columns: IGEPAL CA-360, TritonX, Tween20) across ten-fold dilutions. Linear model (colored line) was fit for observations (colored points) at each detergent percent.

**Supplementary Figure 3.**
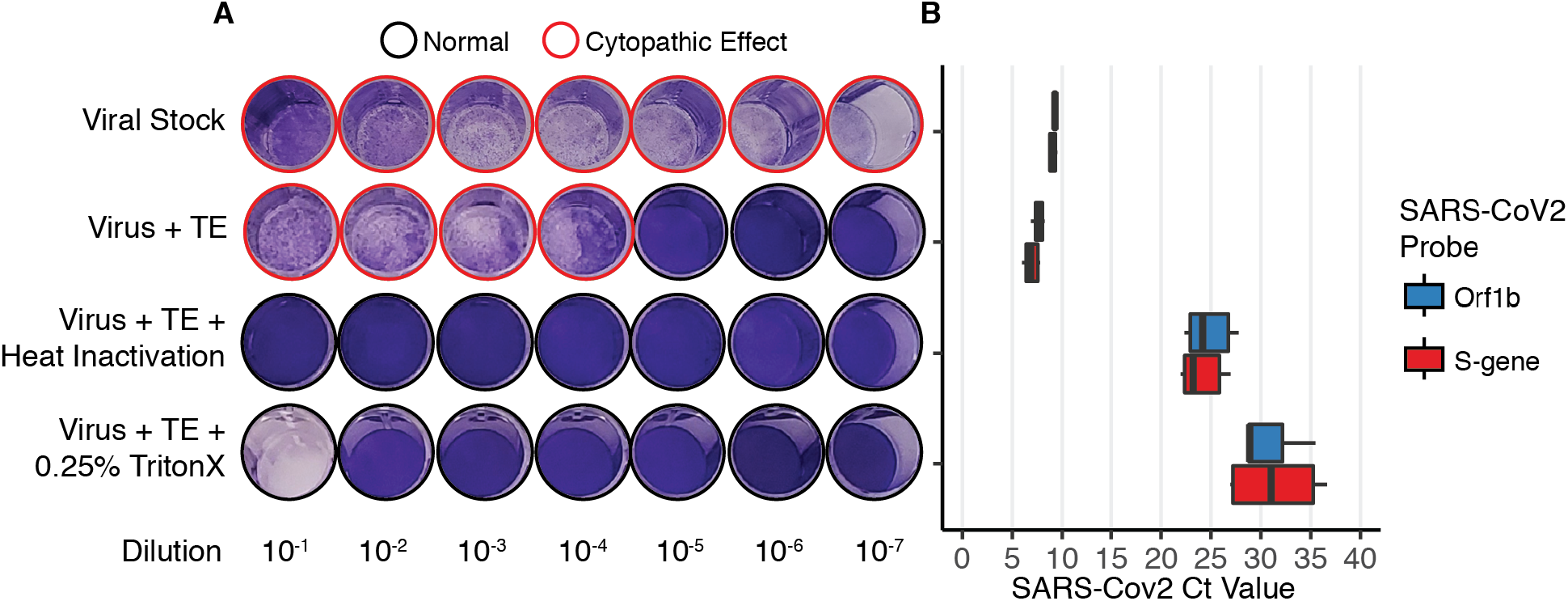
Comparison of heat treatment or detergent to inactivate SARS-CoV-2. (**A**) Crystal violet staining shows the cytopathic effects of SARS-CoV-2 dilutions on Vero cells after incubation with TE, TE + heat inactivation at 65°C for 10 minutes or TE + 0.25% Triton. (**B**) RT-qPCR Ct values of SARS-CoV-2 RNA isolated from Vero cells. Purified viral particles were first incubated with TE, TE + heat inactivation at 65°C for 10 minutes or TE + 0.25% TritonX.

**Supplementary Figure S4.**
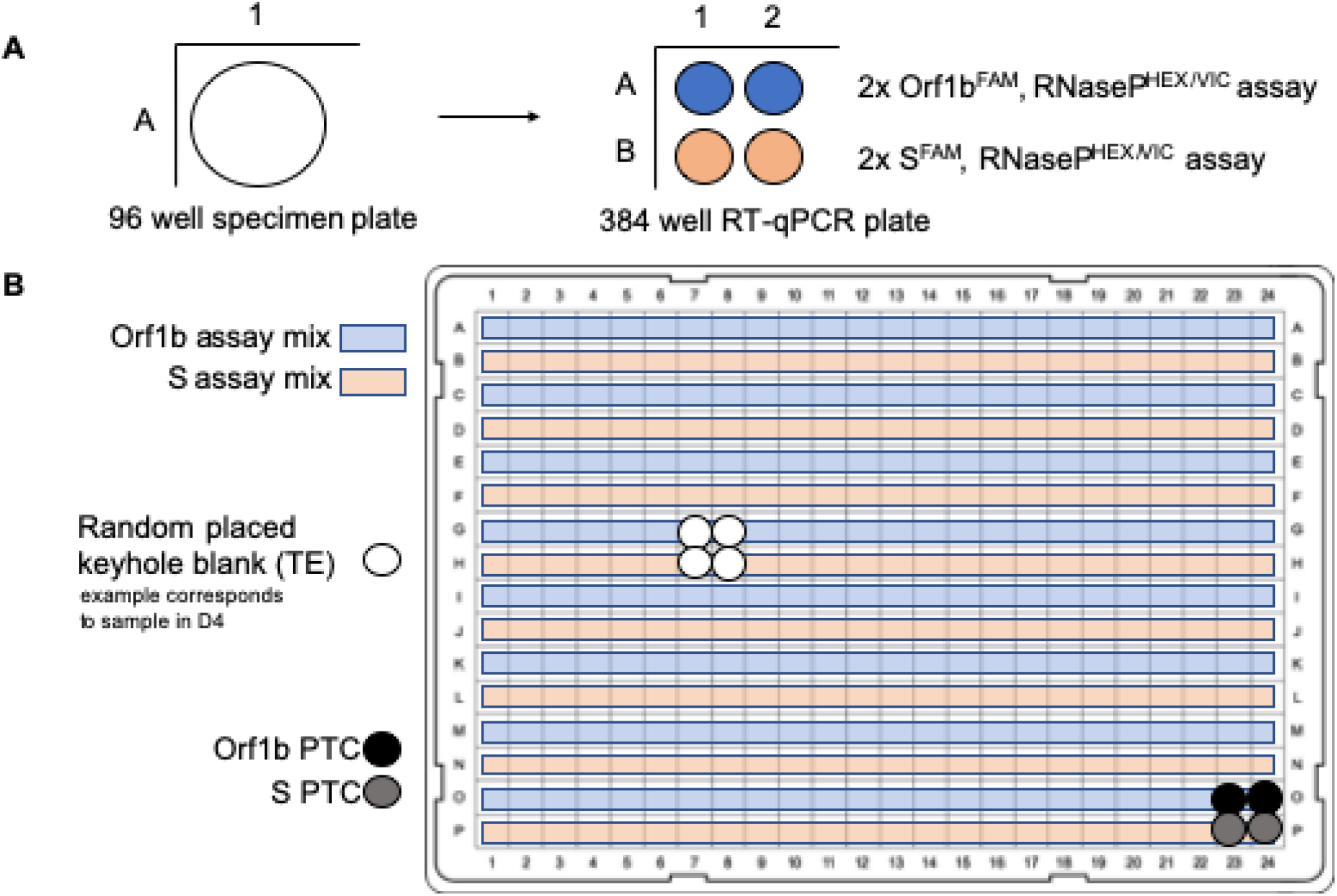
Plate layout of the Northwest Genomics Center SwabExpress RT-qPCR test. (**A**) Each sample is loaded into 4 independent wells on a 96 well plate and tested in duplicate for the Orf1b and S-gene (Spike gene) primer/probes. (**B**) Included on every sample plate is a randomly positioned low-TE keyhole blank (white) and a positive template control (PTC) well containing synthetic SARS-CoV-2 template and human Hap1-RNA (black and grey).

**Supplementary Figure S5.**
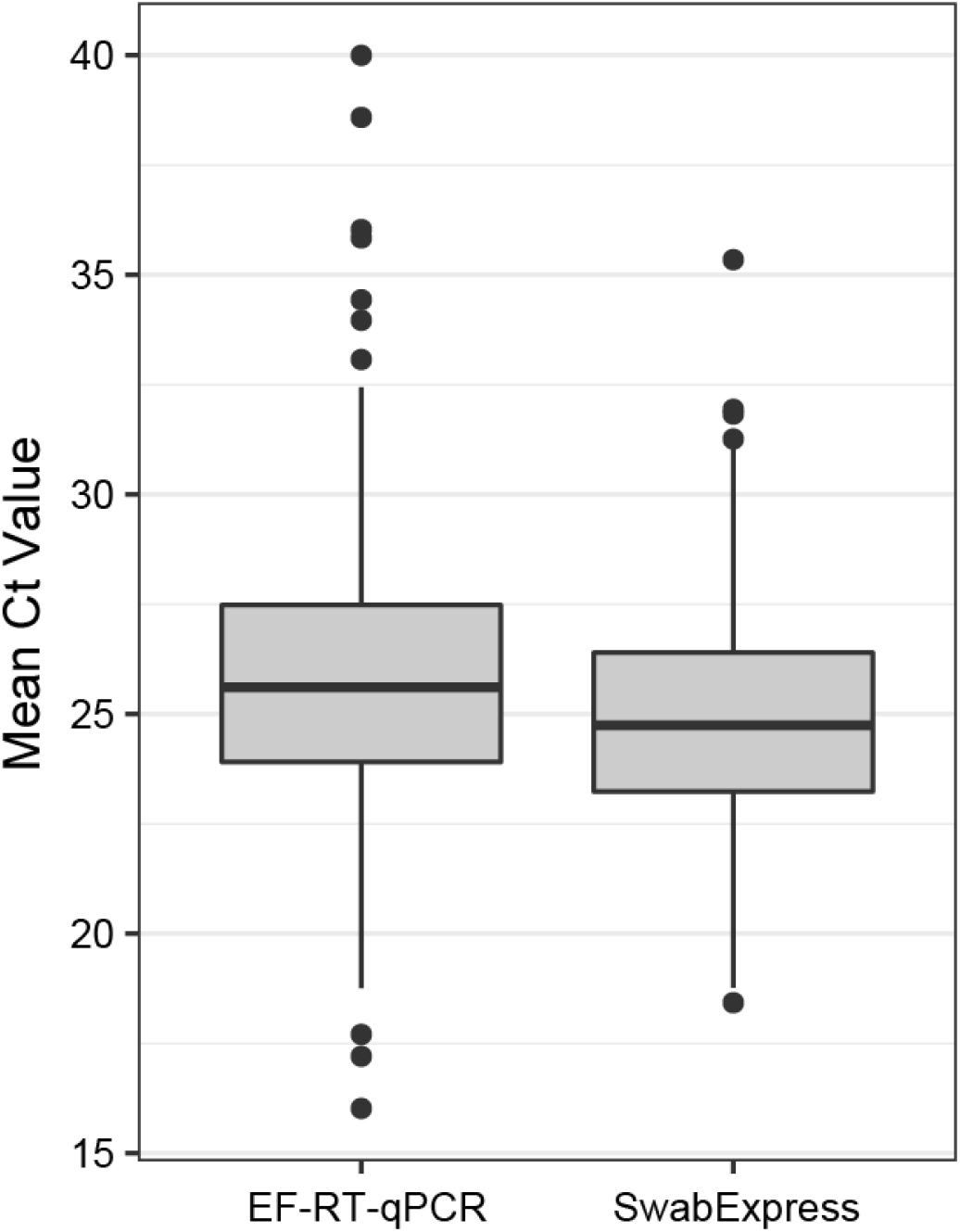
SwabExpress is more sensitive for RNase P than extraction-free RT-qPCR. 1222 samples were tested in parallel by extraction-free RT-qPCR (EF-RT-qPCR) and SwabExpress. Samples run through the SwabExpress protocol, which includes a Proteinase K digestion, had lower mean RNase P Cts than the same samples run on EF-RT-qPCR.

**Supplementary Figure 6.**
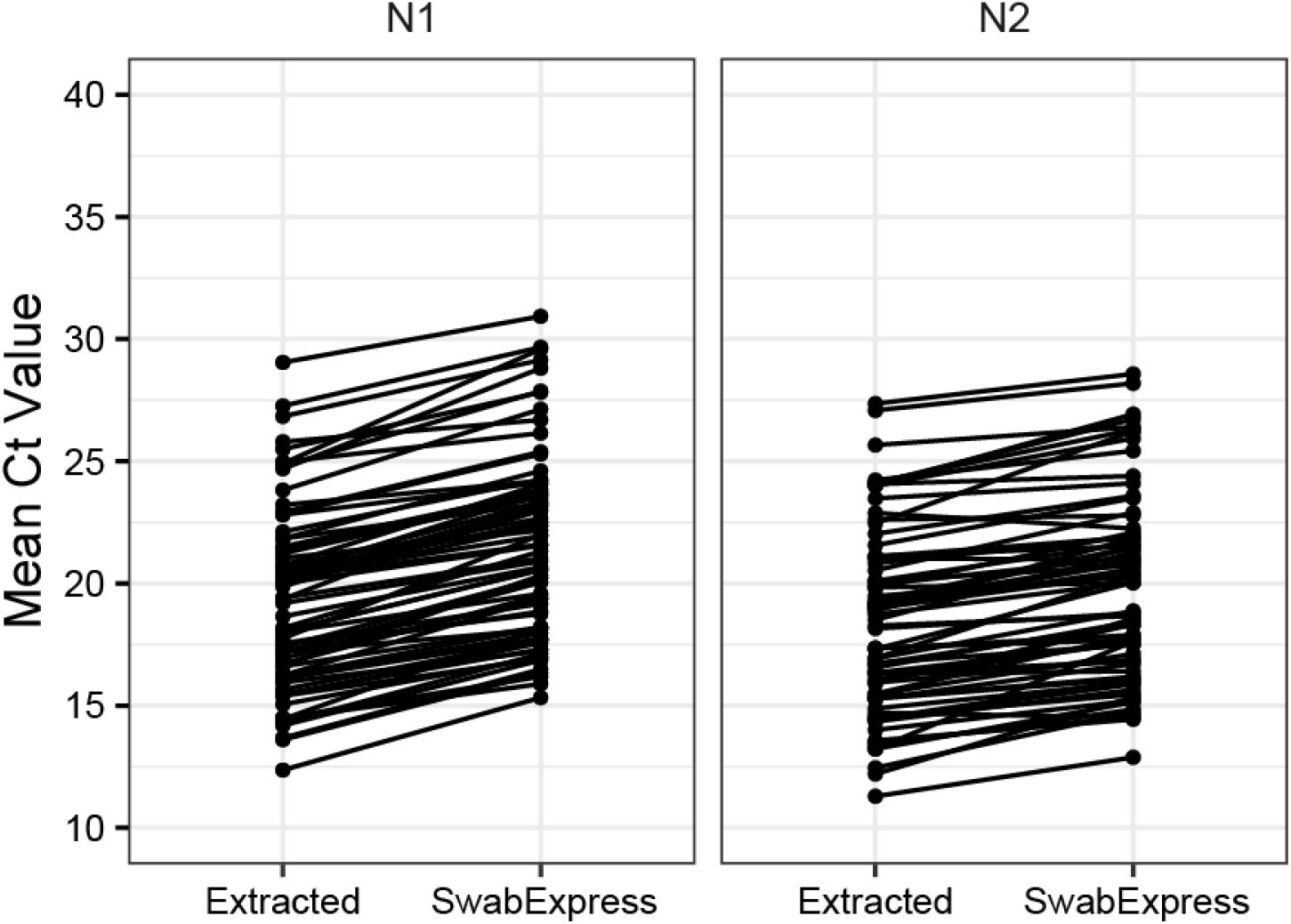
SwabExpress also works with the widely used CDC-N1 and CDC-N2 probesets. Mean crossing threshold for 75 parallel specimens that were extracted or run through SwabExpress for the commonly used SARS-CoV-2 CDC N1 and N2 probe set.

**Table S1A.** Age of study participants.

| Age Groups | Number | % | Average Age<br>(within group) |
| --- | --- | --- | --- |
| Between 18 months - 12 years<br>old | 6 | 17% | 8.7 |
| Between 13 and 18 years old | 8 | 23% | 15.1 |
| Between 18 and 64 years old | 14 | 40% | 40.7 |
| Over 65 years old | 7 | 20% | 68.3 |
| TOTAL | 35 | 100% | 34.9 |
| TOTAL | 35 | 100% | 34.9 |

**Table S1B.** Educational attainment of study participants.

| **Education Levels (of participants 18 & over)** |  |  |  |
| --- | --- | --- | --- |
| **"What is your highest level of education obtained?"** |  |  |  |
| **Response** | **N** | **Participants (%)** | **King County (%)** |
| Less than high school graduate | 0 | 0% | 7% |
| Graduated high school/obtained GED | 1 | 5% | 15% |
| Some college (including vocational training, associate's degree) | 3 | 14% | 18% |
| Bachelor's degree | 11 | 52% | 31% |
| Advanced degree | 6 | 29% | 20% |
| TOTAL | 21 | 100% |  |

**Table S1C.** Household income of study participants.

| **Household Income** |  |  |  |
| --- | --- | --- | --- |
| *"Please choose the range that best represents your household income last year (before taxes). If you are still considered a dependent for tax purposes, choose the range that describes your parent/legal guardian's household income."* |  |  |  |
| Response | N | Study participants (%) | King County (%) |
| Less than or equal to \$25,000 | 0 | 0% | 12% |
| Between \$25,001 to \$50,000 | 0 | 0% | 14% |
| Between \$50,001 to \$75,000 | 1 | 2.9% | 14% |
| Between \$75,001 to \$100,000 | 7 | 20% | 12% |
| Between \$100,001 to \$125,000 | 4 | 11.4% | 19% |
| Between \$125,001 to \$150,000 | 3 | 8.6% | |
| Over \$150,000 | 16 | 45.7% | 29% |
| Don't know | 1 | 2.9% |  |
| Prefer not to say | 3 | 8.6% |  |
| TOTAL | 35 | 100% |  |

**Table S1D.** Race of study participants.

| ***“How would you describe your race? Check all that apply”*** |  |  |  |
| --- | --- | --- | --- |
| Response | N | Study Participants (%) | King County (%) |
| American Indian or Alaska Native | 0 | 0% | <1% |
| Asian | 10 | 24% | 17% |
| Native Hawaiian or Pacific Islander | 0 | 0% | 1% |
| Black or African American | 0 | 0% | 6% |
| White | 30 | 73% | 65% |
| Other | 1 | 3% | 10% |
| Prefer Not to Say | 0 | 0% |  |

**Table S2A.**
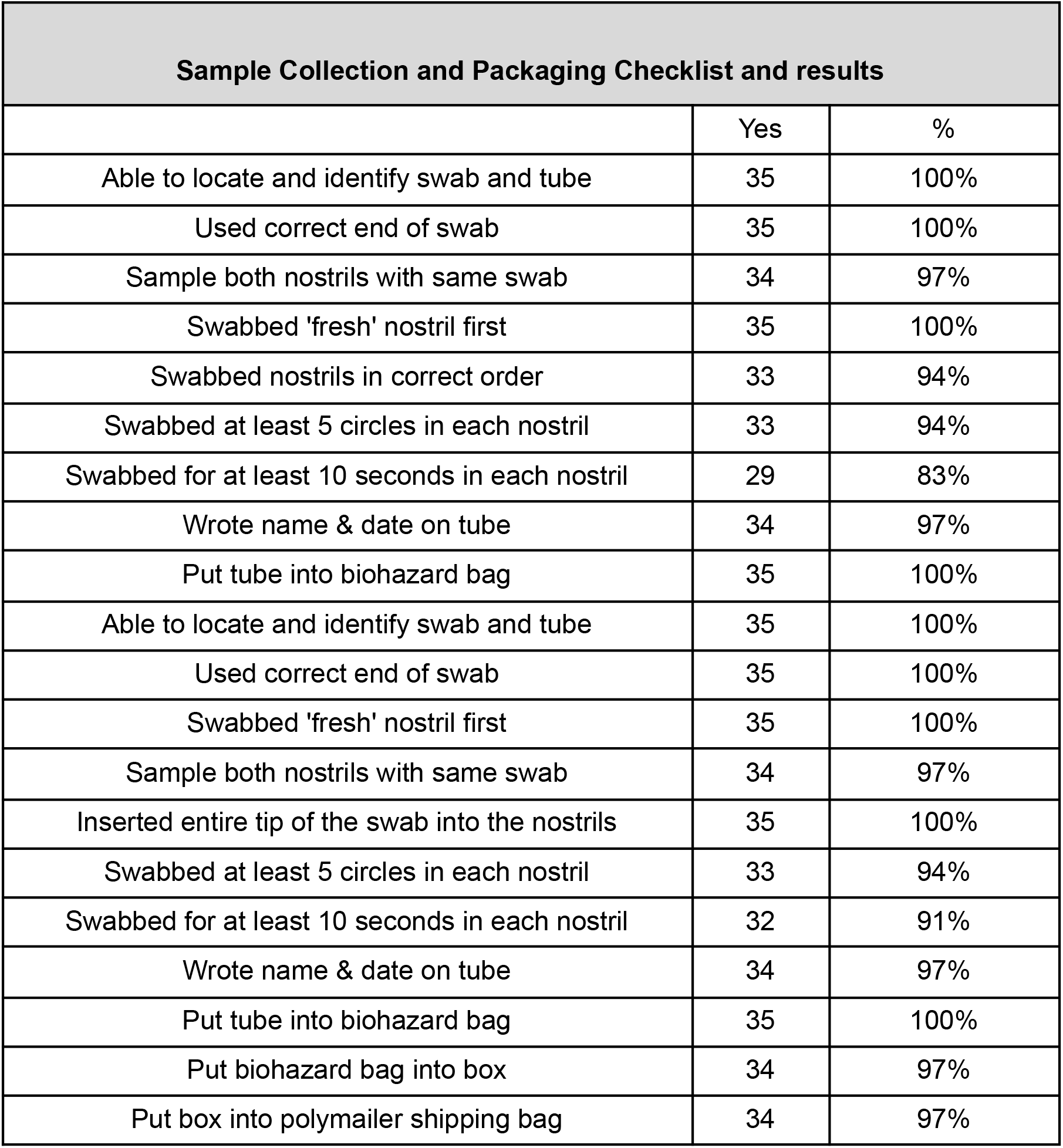
Sample collection and packaging checklist.

**Table S2B.** Observed errors or unwanted outcomes during sample collection or packaging.

| **Did any of the following errors or reactions occur during observation?** |  |  |
| --- | --- | --- |
|  | **N** | **%** |
| Contaminated end of swabs with hand/fingers | 0 | 0% |
| Contaminated end of swabs by setting it down on a table or non-sanitary surface | 0 | 0% |
| Did not put both tubes in biohazard bag | 0 | 0% |
| Did not put biohazard bag in box | 1 | 3% |
| Did not put box in polymailer shipping bag | 1 | 3% |
| Did not write name on tube | 0 | 0% |
| Did not write date on tube | 0 | 0% |
| Nose bleed | 0 | 0% |
| Sneezed during or after swabbing | 4 | 11% |
| Expressed frustration or confusion about one or more of the steps | 3 | 9% |
| Other* | 2 | 6% |

**Table S3.** Summary of SwabExpress limit of detection (LoD) assay results. Lowest concentration with 95% of the replicates detected for each probe is highlighted (in light grey) and bolded.

| Molecules /<br>5uL<br>Reaction | Assay | Percent<br>Detected | Number of<br>replicates | Number<br>of<br>replicates<br>detected | Average<br>Ct | Standard<br>deviation | %Coefficient<br>of variation |
| --- | --- | --- | --- | --- | --- | --- | --- |
| 5 | Spike | 50.0% | 20 | 10 | 36.03 | 0.51 | 1.40% |
| 10 | Spike | 90.0% | 20 | 18 | 34.22 | 0.65 | 1.90% |
| **20** | **Spike** | **95.4%** | **21** | **22** | **34.49** | **0.47** | **1.35%** |
| 40 | Spike | 100% | 24 | 24 | 33.76 | 0.94 | 2.78% |
| 80 | Spike | 100% | 24 | 24 | 31.83 | 0.31 | 0.97% |
| 160 | Spike | 100% | 24 | 24 | 30.36 | 0.23 | 0.76% |
| 320 | Spike | 100% | 24 | 24 | 30.08 | 0.15 | 0.51% |
| 640 | Spike | 100% | 24 | 24 | 28.86 | 0.27 | 0.92% |
| 5 | Orf1b | 85.0% | 20 | 17 | 36.87 | 0.76 | 2.06% |
| **10** | **Orf1b** | **100%** | **20** | **20** | **34.80** | **0.53** | **1.52%** |
| 20 | Orf1b | 100% | 22 | 22 | 34.56 | 0.48 | 1.38% |
| 40 | Orf1b | 100% | 24 | 24 | 33.94 | 0.50 | 1.48% |
| 80 | Orf1b | 100% | 24 | 24 | 32.15 | 0.26 | 0.79% |
| 160 | Orf1b | 100% | 24 | 24 | 31.01 | 0.24 | 0.77% |
| 320 | Orf1b | 100% | 24 | 24 | 30.74 | 0.31 | 1.01% |
| 640 | Orf1b | 100% | 24 | 24 | 29.56 | 0.28 | 0.94% |

**Table S4.**
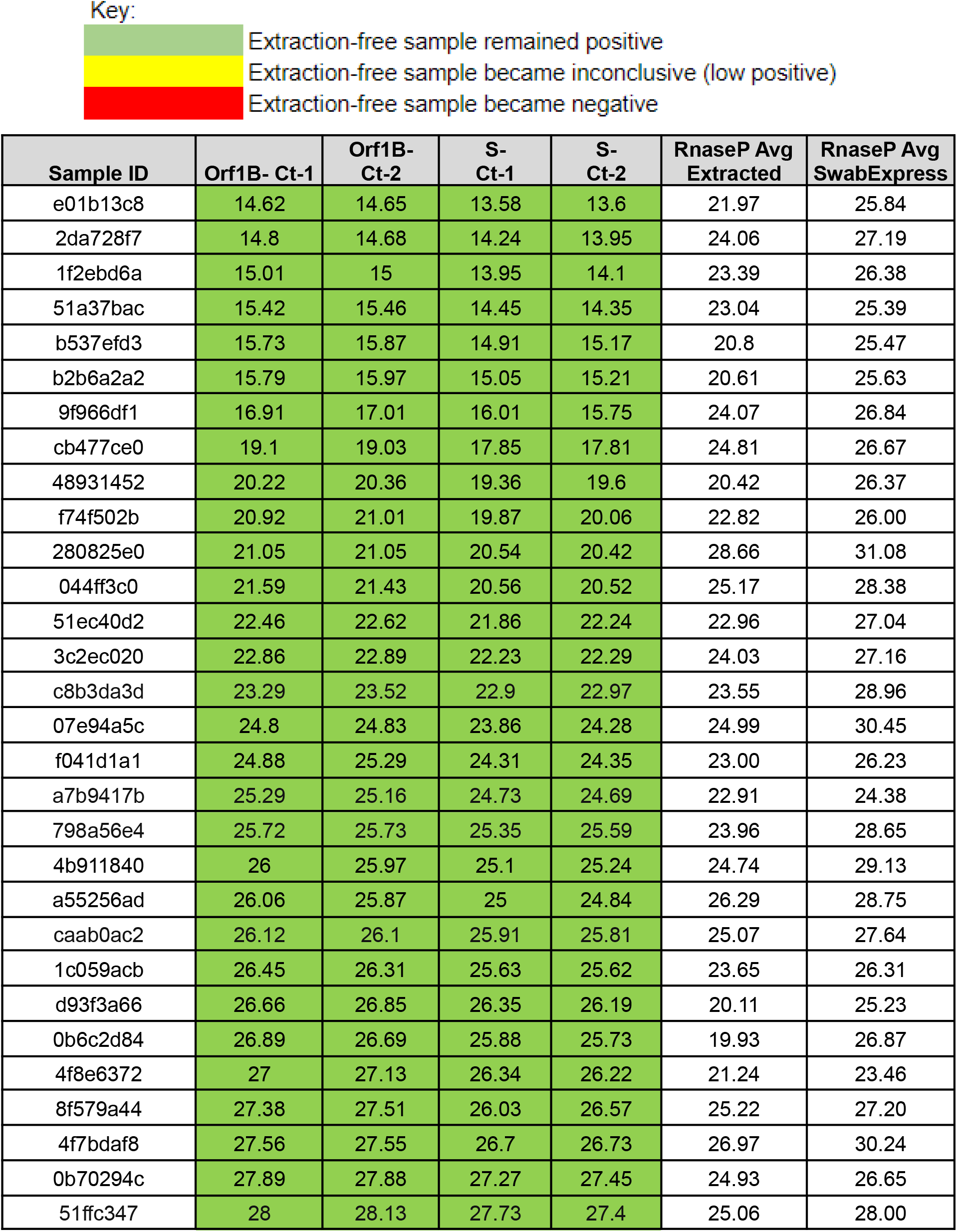
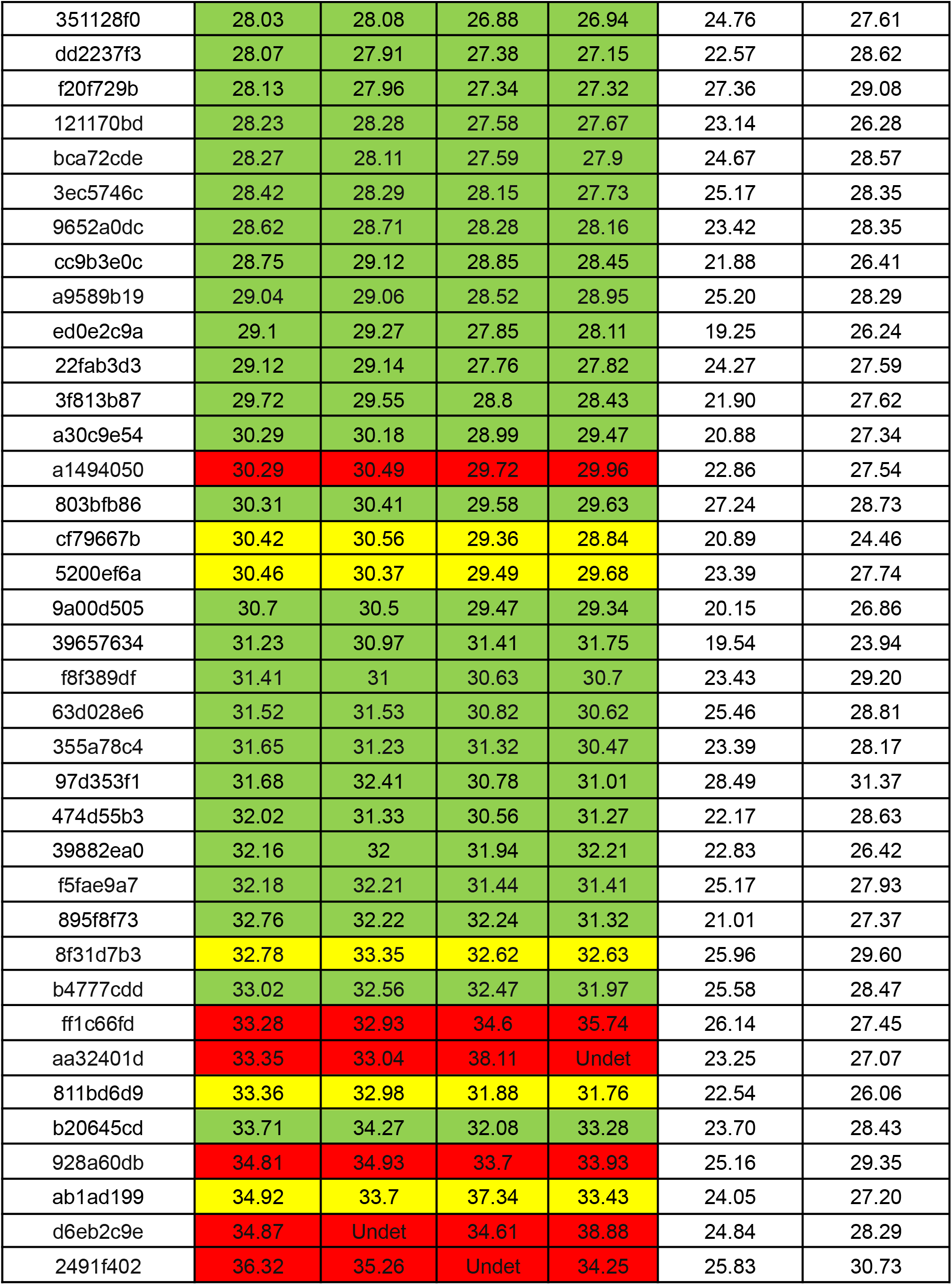
Concordance between extracted nucleic acids and SwabExpress for 67 previously positive specimens. Cts are displayed for extracted specimens and sorted on the first Orf1b Ct. Concordance between extraction and SwabExpress is depicted by color.

**Table S5.** Concordance of 619 AN specimens between KingFisher nucleic acids extraction (KF) and EF-RT-qPCR (EF).

|  | Plate 1<br>COV119 |  | Plate 2<br>RIP107 |  | Plate 3<br>COV120 |  | Plate 4<br>RIP108 |  | Plate 5<br>RIP109 |  | Plate 6<br>RIP112 |  | Plate 7<br>COV179 |  |
| --- | --- | --- | --- | --- | --- | --- | --- | --- | --- | --- | --- | --- | --- | --- |
| **Swab collection type** | Observed |  | At-home |  | Observed |  | At-home |  | At-home |  | At-home |  | Observed , at-home |  |
| **Date** | Oct 26 |  | Oct 26 |  | Oct 26 |  | Oct 27 |  | Oct 27 |  | Oct 28 |  | Nov 8 |  |
| **Operator** | SC | SC | EM | EM | EM | EM | WZ | WZ | WZ | WZ | BB | BB | BB | BB |
| **QuantStudio** | 1 | 2 | 1 | 2 | 1 | 2 | 1 | 2 | 1 | 2 | 2 | 1 | 1 | 2 |
| **Clinical result** | KF | EF | KF | EF | KF | EF | KF | EF | KF | EF | KF | EF | KF | EF |
| **Positive** | 0 | 0 | 4 | 4 | 0 | 0 | 3 | 2 | 0 | 0 | 1 | 1 | 1 | 1 |
| **Low- positive (inconclusive )** | 1 | 0 | 0 | 0 | 0 | 0 | 0 | 2 | 1 | 0 | 0 | 0 | 0 | 0 |
| **Negative** | 87 | 88 | 83 | 83 | 87 | 88 | 87 | 86 | 89 | 90 | 85 | 84 | 89 | 89 |
| **RNase P failures** | 0 | 0 | 0 | 0 | 1 | 0 | 0 | 0 | 0 | 0 | 0 | 1 | 0 | 0 |
| **Total** | 88 | 88 | 87 | 87 | 88 | 88 | 90 | 90 | 90 | 90 | 86 | 86 | 90 | 90 |

**Table S6.** Concordance of prospectively collected swab specimens with or without extraction.

|  |  | **SARS-CoV-2 test with**<br>**KingFisher nucleic acids extraction** |  |  |  |
| --- | --- | --- | --- | --- | --- |
|  |  | positive | inconclusive | negative | test fail |
| **SwabExpress** | positive | 8 | 0 | 0 | 0 |
|  | inconclusive | 1 | 0 | 1 | 0 |
|  | negative | 0 | 2 | 605 | 1 |
|  | test fail | 0 | 0 | 1 | 0 |
| Total |  | 9 | 2 | 607 | 1 |
| positive agreement (9/9) |  | 100% |  |  |  |
| inconclusive agreement (0/2) |  | 0% |  |  |  |
| negative agreement (605/606) |  | 99.8% |  |  |  |
| failed due to poor quality (1/619) |  | 0.16% |  |  |  |

**Table S7.** Comparison of Ct values for SARS-CoV-2 targets from specimens by the SARS-CoV-2 test with KingFisher extraction (KF) and SwabExpress (EF)

| | KF - Orf1b | KF Spike | EF - Orf1b | EF - S gene | KF | EF | ΔCt | ΔCt |
| --- | --- | --- | --- | --- | --- | --- | --- | --- |
| Clinical Sample | Mean ± SD | Mean ± SD | Mean ± SD | Mean ± SD | Clinical Result | Clinical Result | Orf1b | Spike |
| 87359aac | 36.85 * | 38.32 * | undet | undet | incon | neg | N/A | N/A |
| 7bd7b5e1 | 27.41 ± 0.11 | 25.42 ± 0.08 | 29.92 ± 0.00 | 29.05 ± 0.03 | pos | pos | 2.51 | 3.63 |
| d0023767 | 26.07 ± 0.01 | 25.48 ± 0.18 | 27.92 ± 0.11 | 27.15 ± 0.13 | pos | pos | 1.85 | 1.67 |
| 4757675b | 29.62 ± 0.11 | 27.95 ± 0.10 | 30.25 ± 0.02 | 28.61 ± 0.42 | pos | pos | 0.63 | 0.66 |
| 7a22a1f3 | 27.88 ± 0.13 | 27.1 ± 0.21 | 28.67 ± 0.14 | 27.51 ± 0.06 | pos | pos | 0.79 | 0.41 |
| 57f5bb53 | 13.50 ± 0.11 | 12.55 ± 0.45 | 15.66 ± 0.05 | 15.04 ± 0.35 | pos | pos | 2.16 | 2.49 |
| 19344233 | 20.18 ± 0.04 | 19.31 ± 0.23 | 21.66 ± 0.03 | 21.25 ± 0.11 | pos | pos | 1.48 | 1.94 |
| f343051c | 35.22 ± 0.92 | 34.82 ± 0.03 | 36.03 ± 0.36 | undet | pos | incon | 0.81 | NA |
| cc80131a | undet | undet | undet | 39.28 ± 0.60 | neg | incon | NA | NA |
| aef02e4c | 36.6 * | 39.05 * | undet | undet | incon | neg | NA | NA |
| 20e0f036 | 25.44 ± 0.16 | 24.21 ± 0.08 | 29.71 ± 0.29 | 29.50 ± 0.02 | pos | pos | 4.27 | 5.29 |
| 6b496def | 16.92 ± 0.03 | 16.08 ± 0.13 | 20.07 ± 0.00 | 19.61 ± 0.38 | pos | pos | 3.15 | 3.53 |

\* only 1 replicate; pos - Positive; neg - Negative; incon - Inconclusive; undet - Undetermined Ct > 40

**Table S8.** Summary of RNase P detection failure by sample (SwabExpress)

| **RNase P reactions:** | **Number of Specimens in group** |
| --- | --- |
| Failed 1 RNase P reaction | 563 |
| Failed 2 RNase P reactions | 151 |
| Failed 3 RNase P reactions | 38 |
| Failed 4 RNase P reactions | 194 |

**Table S9.**
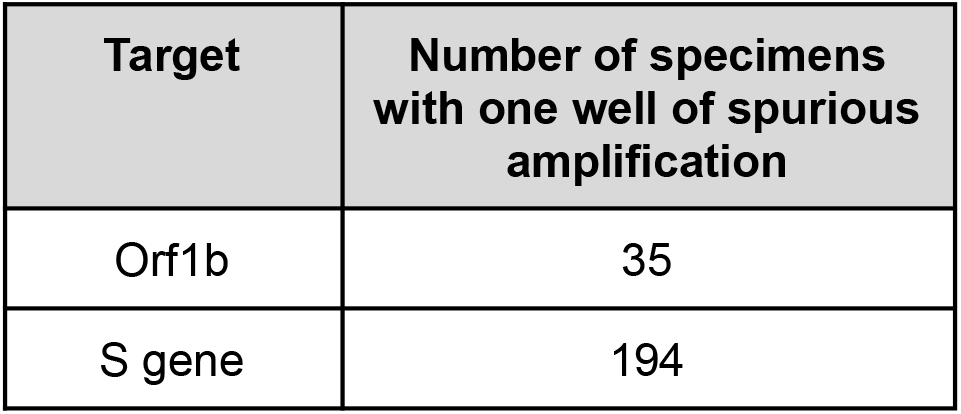
Counts of Spurious (Ct < 30) SARS-CoV-2 amplification from EF-RT-qPCR.

| **Target** | **Number of specimens**<br>**with one well of spurious**<br>**amplification** |
| --- | --- |
| Orf1b | 35 |
| S gene | 194 |

**Table S10.**
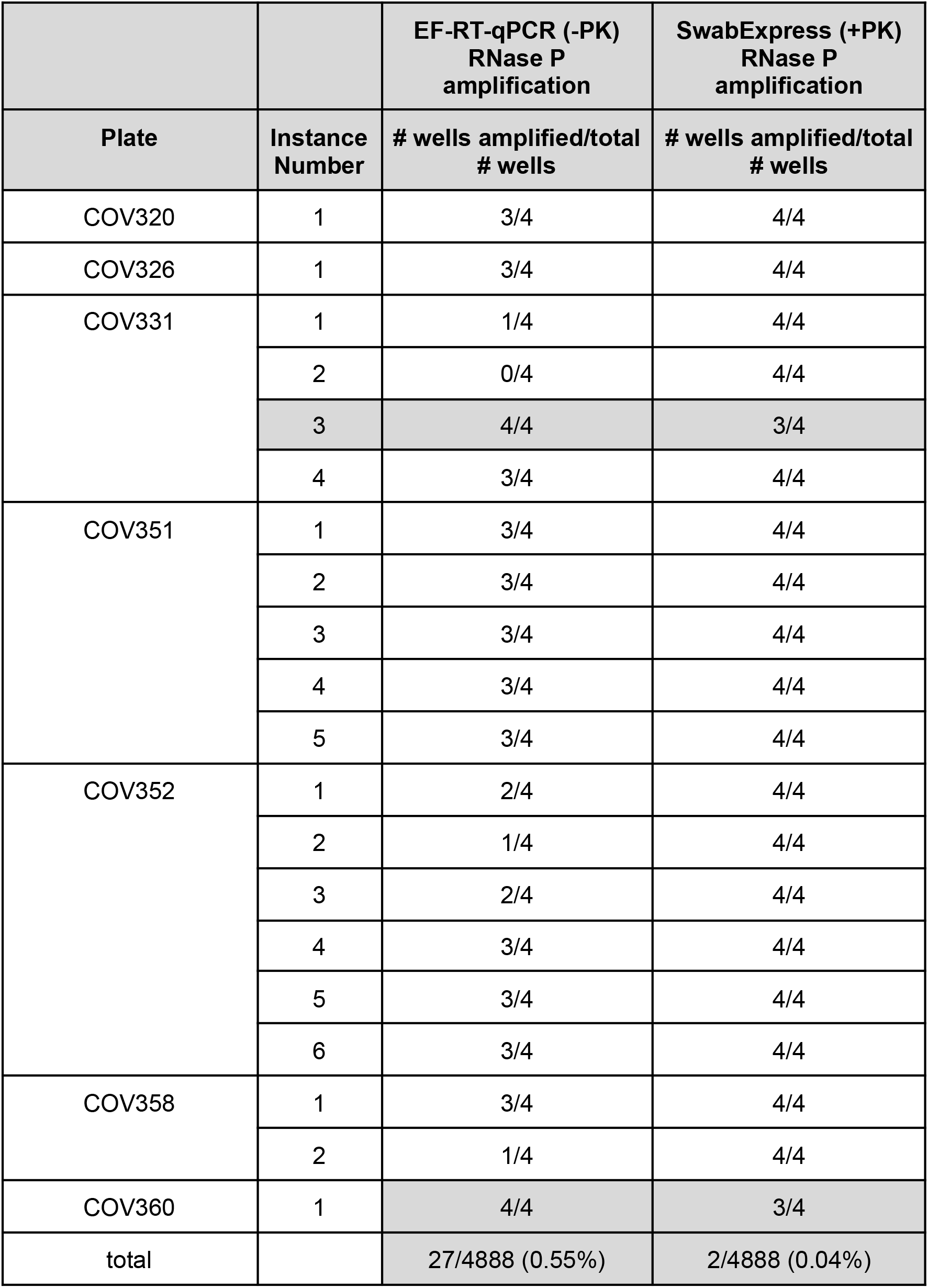
Comparison of RNase P detection failure with and without Proteinase K (PK) digestion.

|  |  | **EF-RT-qPCR (-PK)**<br>**RNase P**<br>**amplification** | **SwabExpress (+PK)**<br>**RNase P**<br>**amplification** |
| --- | --- | --- | --- |
| **Plate** | **Instance**<br>**Number** | **# wells amplified/total**<br>**# wells** | **# wells amplified/total**<br>**# wells** |
| COV320 | 1 | 3/4 | 4/4 |
| COV326 | 1 | 3/4 | 4/4 |
| COV331 | 1 | 1/4 | 4/4 |
|  | 2 | 0/4 | 4/4 |
|  | 3 | 4/4 | 3/4 |
|  | 4 | 3/4 | 4/4 |
| COV351 | 1 | 3/4 | 4/4 |
|  | 2 | 3/4 | 4/4 |
|  | 3 | 3/4 | 4/4 |
|  | 4 | 3/4 | 4/4 |
|  | 5 | 3/4 | 4/4 |
| COV352 | 1 | 2/4 | 4/4 |
|  | 2 | 1/4 | 4/4 |
|  | 3 | 2/4 | 4/4 |
|  | 4 | 3/4 | 4/4 |
|  | 5 | 3/4 | 4/4 |
|  | 6 | 3/4 | 4/4 |
| COV358 | 1 | 3/4 | 4/4 |
|  | 2 | 1/4 | 4/4 |
| COV360 | 1 | 4/4 | 3/4 |
| total |  | 27/4888 (0.55%) | 2/4888 (0.04%) |

**Table S11.** Concordance of clinical results for specimens with total nucleic acids extraction (Kingfisher Flex), heat treatment (EF-RT-qPCR) or Proteinase K digestion plus heat treatment (SwabExpress)

|  | Sample Prep Method |  |  |
| --- | --- | --- | --- |
| Qualitative Result | KingFisher Flex | EF-RT-qPCR | SwabExpress |
| # Positive | 23 | 24 | 28 |
| # Inconclusive | 3 | 3 | 0 |
| # Negative | 4 | 3 | 2 |

**Table S12.**
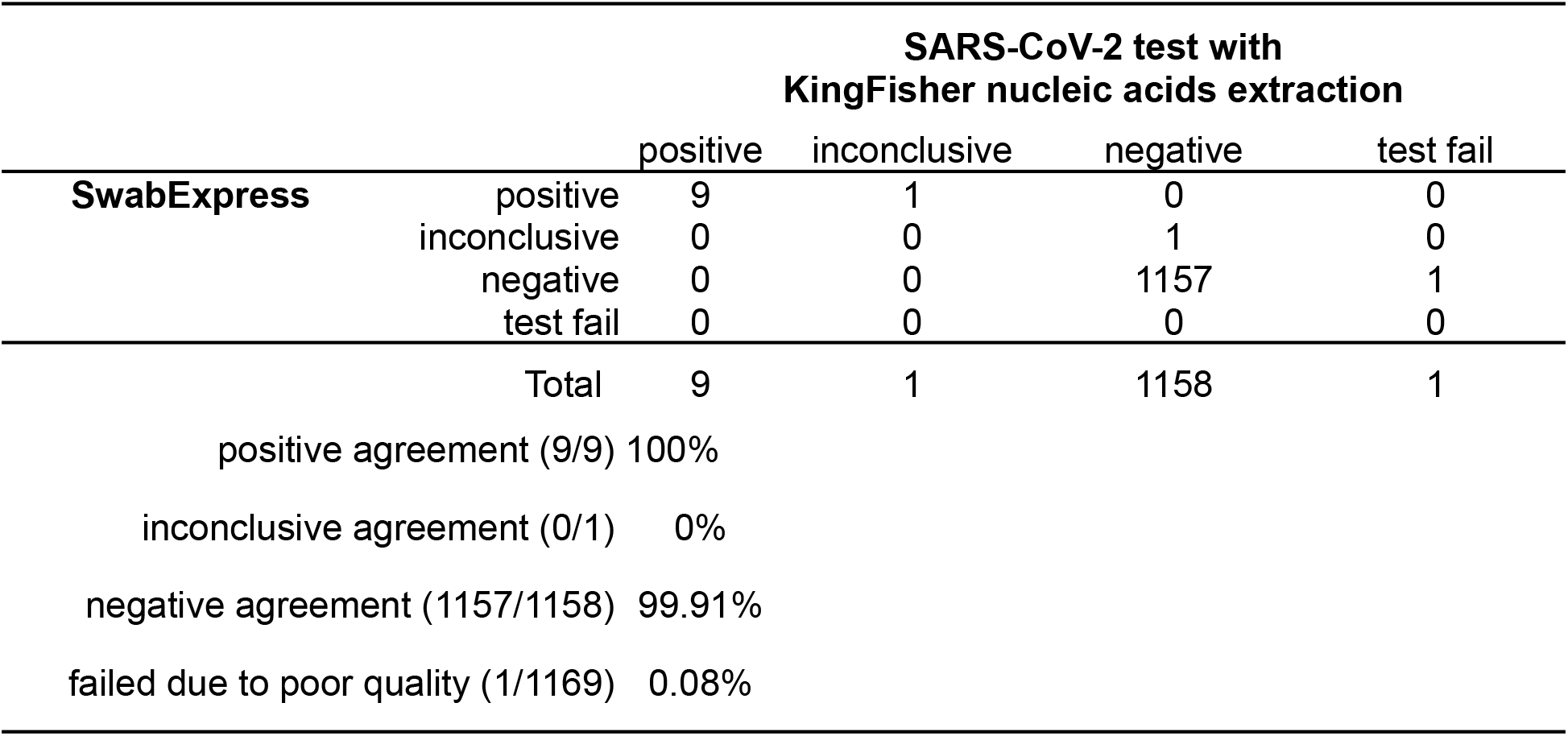
Concordance of prospectively collected swab specimens from participants with or without extraction.

**Table 13.** Mean ± SD of 75 positive specimens extracted on the Roche Magna Pure 96 (MP96) or processed SwabExpress (SE) protocol and amplified using the N1 and N2 CDC probe sets.

| Probe | MP96<br>Mean ± SD | SwabExpress<br>Mean ± SD | ΔCt<br>[SE-MP96] |
| --- | --- | --- | --- |
| N1 | 19.22 ± 3.67 | 21.79 ± 4.33 | 2.57 |
| N2 | 18.31 ± 3.73 | 19.80 ± 3.72 | 1.49 |

**Table S14.** SwabExpress per sample cost breakdown. Values displayed in US dollars.

| Item | SwabExpress |
| --- | --- |
| US Cotton #3 Swab | \$0.29 |
| Transport Tube | \$0.61 |
| Barcode Stickers | \$0.50 |
| Low TE | \$0.05 |
| Proteinase K | \$0.14 |
| RT-qPCR Reagents | \$2.67 |
| Lab Consumables | \$1.72 |

